# RNA-Sequencing of Long-Term Label-Retaining Colon Cancer Stem Cells Identifies Novel Regulators of Quiescence

**DOI:** 10.1101/2021.02.02.429354

**Authors:** Joseph L. Regan, Dirk Schumacher, Stephanie Staudte, Andreas Steffen, Ralf Lesche, Joern Toedling, Thibaud Jourdan, Johannes Haybaeck, Dominik Mumberg, David Henderson, Balázs Győrffy, Christian R.A. Regenbrecht, Ulrich Keilholz, Reinhold Schäfer, Martin Lange

## Abstract

Recent data suggests that colon tumors contain a subpopulation of therapy resistant quiescent cancer stem cells (qCSCs) that are the source of relapse following treatment. Here, using colon cancer patient-derived organoids (PDOs) and xenograft (PDX) models, we identify a rare population of long-term label-retaining (PKH26^Positive^) qCSCs that can re-enter the cell cycle to generate new tumors. RNA-sequencing analyses demonstrated that these cells are enriched for stem cell associated gene sets such as Wnt and hedgehog signaling, epithelial-to-mesenchymal transition (EMT), embryonic development, tissue development and p53 pathway but have downregulated expression of genes associated with cell cycle, transcription, biosynthesis and metabolism. Furthermore, qCSCs are enriched for p53 interacting negative regulators of cell cycle, including *AKAP12, CD82, CDKN1A, FHL2, GPX3, KIAA0247, LCN2, TFF2, UNC5B* and *ZMAT3*, that we show are indicators of poor prognosis and may be targeted for qCSC abolition. Interestingly, CD82, KIAA0247 and UNC5B proteins localize to the cell surface and may therefore be potential markers for the prospective isolation of qCSCs. These data support the temporal inhibition of p53 signaling for the elimination of qCSCs and prevention of relapse in colorectal cancer.

## INTRODUCTION

Molecular and functional intra-tumoral heterogeneity contribute to differences in treatment outcomes between colorectal cancer (CRC) patients with similar mutational profiles^1^. Studies of functional heterogeneity, as defined by phenotypic differences between cells, suggest that cancer stem cells (CSCs) are responsible for tumor growth, metastasis and therapy resistance^2–7^. CSCs share many of the characteristics of normal tissue stem cells, including unlimited self-renewal, the ability to generate differentiated daughter cells and chemoresistance^8,9^.

The normal intestine is maintained by highly clonogenic crypt base LGR5^Positive^ stem cells and also contains a population of rare quiescent (G0 phase) stem cells that act as a clonogenic reserve capable of re-entering the cell cycle upon perturbation of tissue homeostasis, e.g. after injury leading to loss of the cycling crypt base stem cells^10–15^. Cancer often recapitulates the cellular hierarchy of the tissue in which it arises, and recent evidence suggests that many tumor types contain rare slow cycling / qCSCs^16–27^. Conventional chemotherapies and radiotherapies target proliferating cells and require active cycling for induction of apoptosis^1^. In addition, cellular quiescence has been shown to facilitate immune evasion^28^. Thus, non-dividing qCSCs may escape conventional therapeutic strategies and represent the source of disease relapse after treatment^2,29–31^.

Cell cycle activation in qCSCs has been proposed as a therapeutic strategy to sensitize qCSCs to treatment and lead to long-term disease-free survival without relapse^29,30^. However, the molecular profiling of qCSCs for the identification of novel cell cycle regulators that do not also perturb cellular homeostasis in healthy tissues has been limited by both the rarity of qCSCs and the small number of suitable experimental assays available for their detection. PDOs echo the morphological, differentiation, intratumor mutational and drug sensitivity status of the original tumor^32,33^ and thus provide an excellent model for the prospective isolation and profiling of qCSCs.

Strategies for the identification of quiescent cells employ pulse-chase approaches, including label retention (e.g. BrdU, PKH26, CFSE), wherein dividing cells lose the label and quiescent or slow cycling cells retain the label for an extended period of time, or the dilution of histone 2B-GFP (H2B-GFP)^34^. In contrast to the H2B-GFP approach^35^, which can identify transient quiescent cells, label retention allows for the identification of cells that remain quiescent from the early stages of tumorigenesis. This is important since cells selectively surviving chemotherapy have been shown to be the same cells that are quiescent/slow cycling in untreated tumors and not cells that became quiescent upon drug treatment^36^. Such label-retaining cells (LRCs) have previously been reported in colon cancer cell lines, xenografts and, more recently, in PDOs^6,36–38^.

However, to date, the transcriptomic profiling of qCSCs in CRC patients has been limited to microarray analyses of transiently slow-cycling H2B-GFP^Positive^ cells from a single CRC patient by Puig *et al*. (2018)^35^ and of PKH26^Positive^ LRCs from two colon cancer patient-derived (via spheroid culture) xenograft models by Francescangeli *et al*. (2020)^36^. In addition, the LRCs reported in the latter study were not functionally tested for proliferative or self-renewal capacity prior to molecular profiling and instead relied on expression of CD133 as evidence of a stem cell phenotype. CD133 expression is not restricted to stem cells of the intestine and is expressed on both CSC and differentiated tumor cells^39–42^.

Here, we report the identification and first whole-transcriptome RNA-sequencing analyses of label-retaining qCSCs in a panel of PDOs encompassing primary colon tumors and metastases. These cells maintain a large proliferative capacity, persist long term *in vivo* and display the molecular hallmarks of quiescent tissue stem cells^43^, including enrichment for p53 pathway and developmental gene sets alongside downregulation of cell cycle, transcription, biosynthesis and metabolism genes. In addition, we show that qCSCs are enriched for p53 interacting negative regulators of cell cycle that we propose may be targeted for cell cycle activation and the elimination of qCSCs. These data provide a valuable resource for the development of novel therapeutic strategies geared toward the elimination of minimal residual disease and the prevention of relapse.

## RESULTS

### Colon cancer PDOs contain rare label-retaining qCSCs that persist long term *in vivo*

To determine whether PDOs contain non-cycling LRCs, we performed an initial 72 h pulse chase experiment using CM-DiL dye. PDOs were established as previously described^44,45^, processed to single cells, uniformly labelled with CM-DiL dye and seeded in Matrigel culture. CM-DiL is diluted with each cell division, halving its fluorescence between each daughter cell until it becomes undetectable. Non-cycling cells can thus be identified by their label retention. After 72 h the majority of PDO cells had lost the CM-DiL dye but some PDOs contained non-cycling LRCs (Figure 1A). To determine the frequency of these non-cycling (G0) cells we performed EdU cell cycle analysis on a panel of colon cancer PDOs (Table S1). This analysis demonstrated that PDOs contain non-cycling cells that do not proliferate and remain in G0 within a 72 h period (Figure 1B and C).

**Figure 1.**
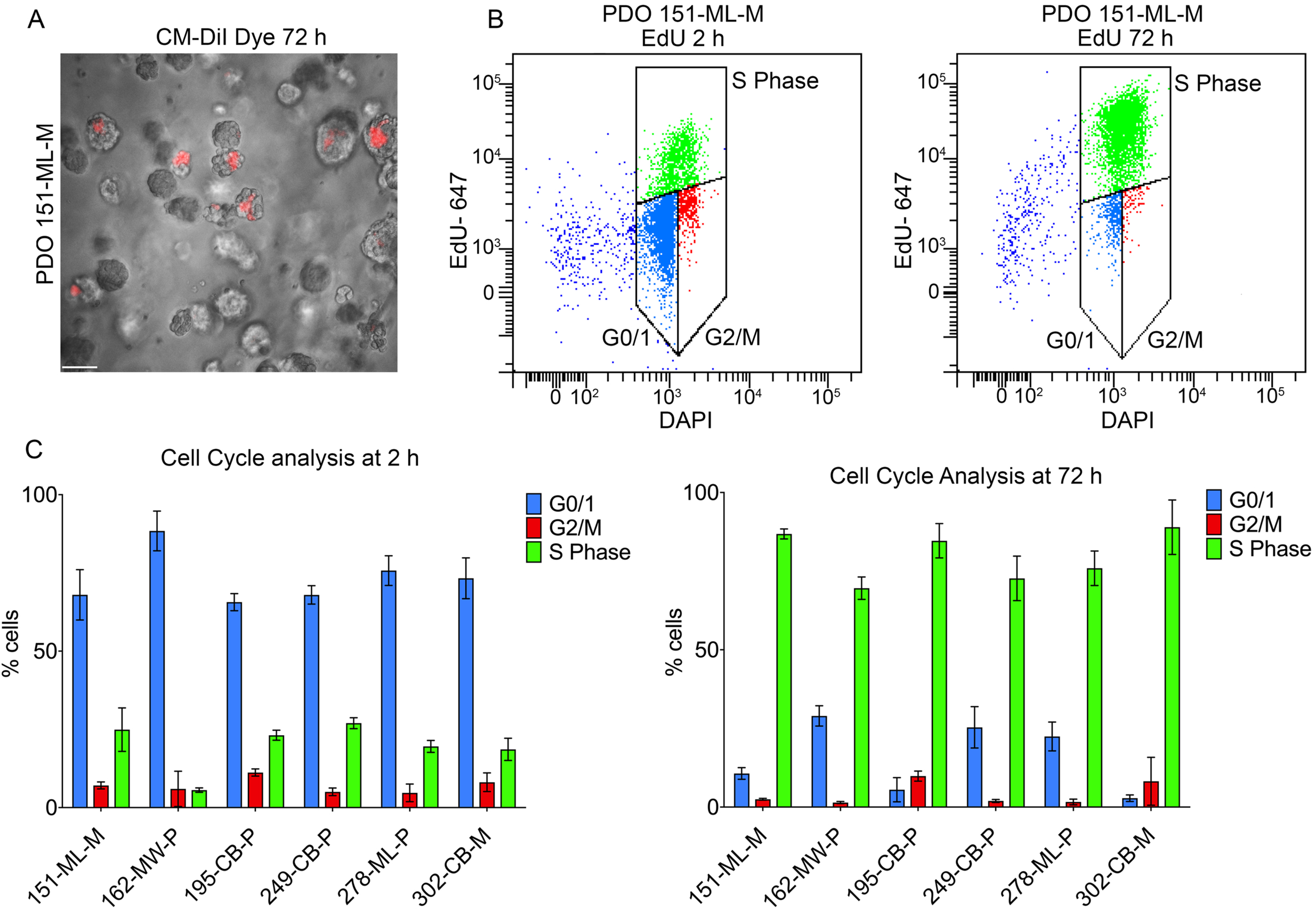
Colon cancer PDOs contain a subpopulation of non-cycling cells. (A) Phase contrast image of colon cancer PDOs labelled with cell-tracker dye CM-DiL after 72 h (Bar = 75 μm) (see also Table S1). (B) Representative FACS plots of EdU cell cycle analysis of 151-ML-M PDO cells at 2 h (left hand side) and 72 h (right hand side) after labelling. (C) Percentage of cells (±SD) in G0/1, G2/M and S Phase at 2 h and 72 h post EdU labelling in PDO models 151-ML-M, 162-MW-P, 195-CB-P, 249-CB-P, 278-ML-P and 302-CB-M (data from three independent experiments).

To determine the long-term proliferative capacity of these non-cycling cells, we labelled cells with the lipophilic fluorescent dye PKH26. Unlike CM-DiL, which is suitable for short term label retention studies, PKH26 labelling can be used to identify non-cycling cells for up to six months (*in vitro* and *in vivo*)^46,47^. PDOs were dissociated to single cells, labelled with PKH26 and replated in Matrigel culture. After 12 days PDOs were re-processed to single cells and analyzed by fluorescence assisted cell sorting (FACS). These data demonstrated that PDOs contain rare, non-cycling, long-term LRCs (Figure 2A and B). Crucially, FACS isolation and replating of PKH26^Positive^ DAPI^Negative^ (live) cells from 12 day cultures demonstrated that they are not label-retaining due to terminal differentiation or senescence but can re-enter the cell cycle to generate organoids and have a large proliferative capacity (Figure 2C – F). In addition, non-adherent spheroid formation assays, the gold standard assay for testing stem cell function *in vitro*^48,49^, showed that PKH26^Positive^ cells are enriched for self-renewing CSCs (Figure 2G).

**Figure 2.**
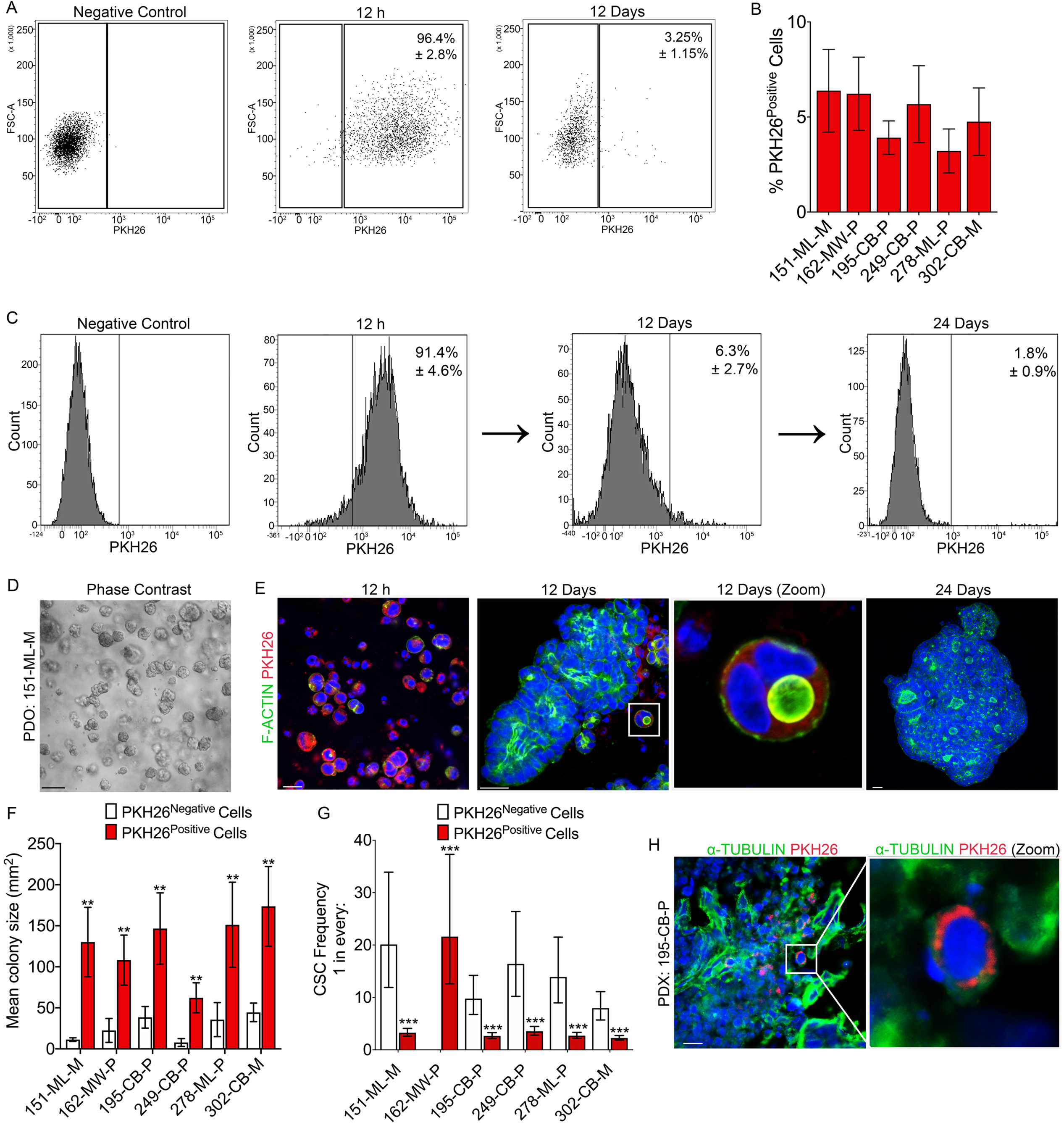
Non-cycling PDO cells are quiescent CSCs that can re-enter cell cycle and persist long-term *in vivo*. (A) Representative FACS plot of PKH26 labelled 278-ML-P PDO cells after 12 h (middle panel) and 12 days (right side panel) compared to non-labelled control (left side panel). (B) Frequency (±SD) of PKH26^Positive^ LRCs in PDO models after 12 days (data from 5 independent experiments). (C) FACS histograms demonstrating frequency of PKH26^Positive^ cells in 151-ML-M PDOs at 12 h (left side panel) and 12 days (middle panel) after staining and 24 days (right side panel) after FACS isolation and serial replating of PKH26^Positive^ cells from 12 day cultures. (D) Phase contrast of unlabeled PDOs (negative control) (Bar = 100 μm) and (E) immunofluorescence images of PKH26 labelled PDOs at 12 h and 12 days (left and middle panels) and 24 days after FACS isolation and serial re-plating of PKH26^Positive^ LRCs from 12 day cultures (right side panel). Cells are stained for F-ACTIN (green) and nuclei are counterstained with DAPI (blue) (Bars = 20 μm). (F) Mean colony size (±SD) of PKH26^Negative^ and PKH26^Positive^ cell derived PDOs in Matrigel culture. Data from three independent experiments. **p-value: < 0.01 (t test). (G) Limiting dilution spheroid formation assay of PKH26^Negative^ and PKH26^Positive^ cells. Data from three independent experiments. The p-values for pairwise tests of differences in CSC frequencies between PKH26^Negative^ and PKH26^Positive^ cells in 151-ML-M, 162-MW-P, 195-CB-P, 249-CB-P, 278-ML-P and 302-CB-M tumors are 1.27 × 10^−13^, 1.87 × 10^−5^, 6.42 × 10^−11^, 1.12 × 10^−10^, 3.5 × 10^−14^, 6.14 × 10^−12^, respectively. (H) Immunofluorescence image of a frozen PDX section derived from 1,000 PKH26 labelled 195-CB-P PDO cells 80 days post transplantation. Magnified region indicates a long-term label-retaining PKH26^Positive^ cell. Cells are stained for *α*-tubulin (green) and nuclei are counterstained with DAPI (blue) (Bar = 100μm).

In order to test whether these cells also persisted long-term *in vivo* we generated xenografts by transplanting PKH26 labelled cells. Long-term tracking of LRCs in xenografts requires the slow growth of the tumor. Cells were therefore transplanted at a low cell number based on knowledge of tumor growth rates from previous limiting dilution xenotransplantation assays, in which xenografts were generated from 1,000 PDO cells^44^.

Unlabeled cells, lacking the burden of carrying a fluorescent dye may be at a competitive advantage over labelled cells. Therefore, immediately prior to transplantation, PKH26 labelled cells were processed by FACS to exclude unlabelled cells and thus ensure that only live (DAPI^Negative^) PKH26 labelled cells would give rise to tumors. Significantly, analysis of xenograft tissue demonstrated the presence of PKH26^Positive^ LRCs for up to 80 days after transplantation (Figure 2H). Previous studies have observed quiescence to be a transient state^35^. However, these data demonstrate that quiescence can be stable and persist long-term from the initial stages of tumor development.

### RNA-sequencing of PKH26^Positive^ cells reveals a molecular signature of qCSCs

To generate a molecular profile of qCSCs we carried out RNA-sequencing analyses of PKH26^Negative^ (cycling) and PKH26^Positive^ (non-cycling) qCSCs isolated from a panel of six different PDO models (Table S1) after 12 days in Matrigel culture. These data demonstrated that PKH26^Positive^ qCSCs are enriched for stem cell associated gene sets, such as embryonic development, organ development, placenta, nervous system development, EMT, Wnt and hedgehog signaling (Figure 3A).

**Figure 3.**
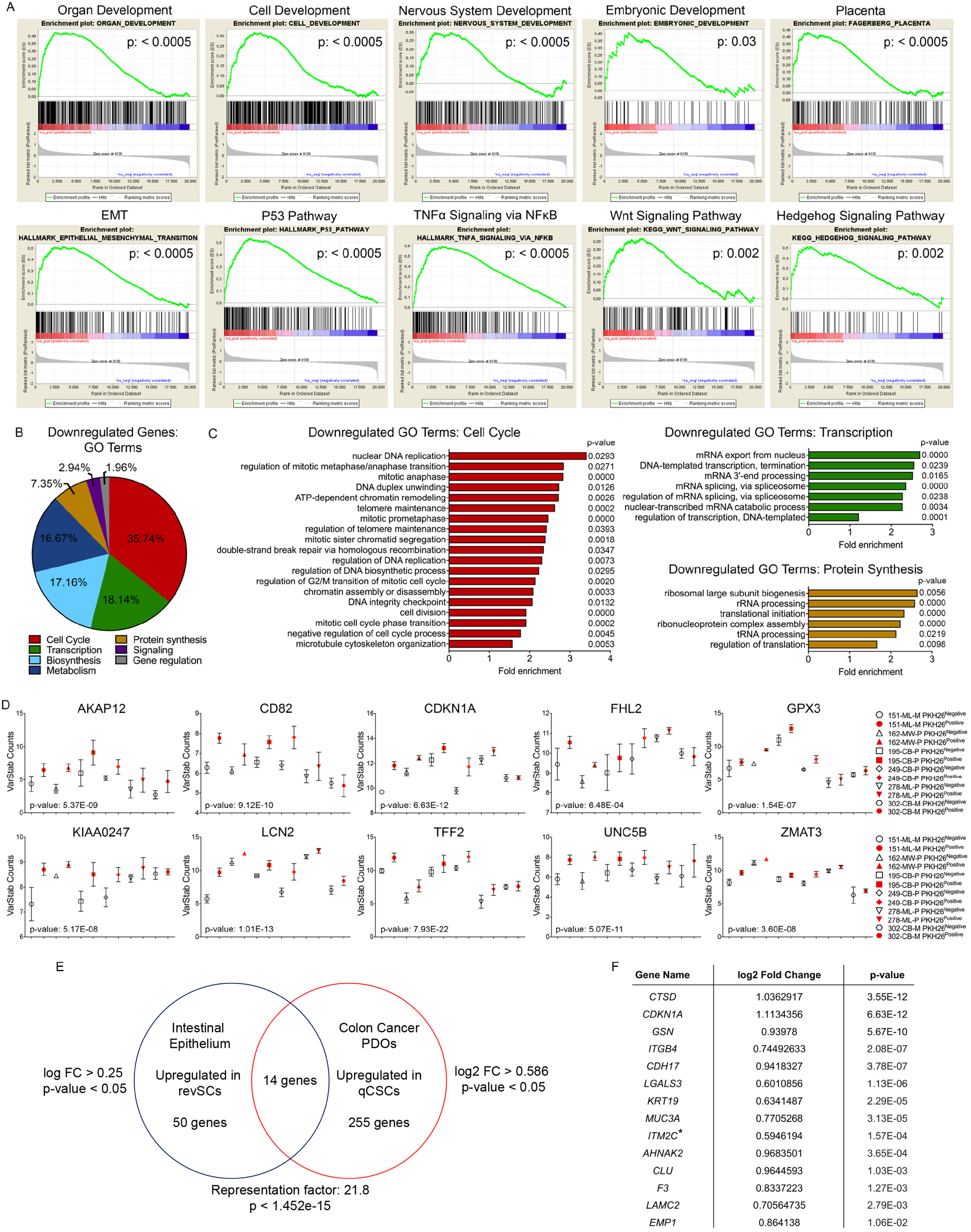
PKH26^Positive^ qCSCs are enriched for stem cell associated gene sets and p53-interacting negative regulators of proliferation and have downregulated expression of cell cycle genes. (A) RNA sequencing generated gene set enrichment analysis for organ development (nominal p-value = < 0.0005), cell development (nominal p-value = < 0.0005), nervous system development (nominal p-value = < 0.0005), embryonic development (nominal p-value = 0.03), placenta (nominal p-value = < 0.0005), epithelial mesenchymal transition (nominal p-value = < 0.0005), p53 pathway (nominal p-value = < 0.0005), TNFa signaling via NFkB (nominal p-value = < 0.0005), Wnt signaling pathway (nominal p-value = 0.002) and hedgehog signaling pathway (nominal p-value = 0.002) in 12 day PKH26^Positive^ LRCs (compared to PKH26^Negative^ cells) from PDO models 151-ML-M, 162-MW-P, 195-CB-P, 249-CB-P, 278-ML-P and 302-CB-M (n = 4 separate cell preparations). (B) Gene ontology (GO) groups downregulated in PKH26^Positive^ LRCs. (C) Cell Cycle, transcription and protein synthesis GO terms downregulated in PKH26^Positive^ LRCs. (D) RNA sequencing generated normalized counts for negative cell cycle regulator and p53 target genes *AKAP12, CD82, CDKN1A, FHL2, GPX3, KIAA0247, LCN2, TFF2, UNC5B* and *ZMAT3* in PKH26^Negative^ and PKH26^Positive^ cells. (E) Venn diagram shows the number of upregulated RNA-sequencing generated transcripts identified in intestinal revSCs (50 genes; log fold change > 0.25, p-value < 0.05) by Ayyaz *et al*. (2019)^15^ and in PKH26^Positive^ qCSCs (255 genes; log2 fold change > 0.586, p-value < 0.05) and upregulated in both revSCs and PKH26^Positive^ qCSCs (14 genes; representation factor 21.8, p-value < 1.452e-15). The representation factor is the number of overlapping genes divided by the expected number of overlapping genes drawn from two independent groups. A representation factor > 1 indicates more overlap than expected of two independent groups. (F) Table shows the 14 genes upregulated in both revSCs and PKH26^Positive^ qCSCs. *ITM2C is a paralog of revSC enriched Itm2b. (See also Figure S1).

At the same time as showing enrichment for genes associated with growth and development, PKH26^Positive^ qCSCs have downregulated cell cycle, transcription, protein synthesis, metabolism and biosynthesis genes (Figure 3B and C). These data are in agreement with the transcriptional profiles of slow cycling / qCSCs reported in previous studies^35–37^ and demonstrate a common molecular signature of qCSCs.

The normal intestine also contains quiescent stem cells that can regenerate the damaged intestine upon loss of crypt base stem cells following injury, although whether cellular plasticity or distinct cell types are responsible for this remains unclear. Bmi1^50^, Hopx^51^, Lrig1^52^ and Tert^53^ have previously been reported as markers of quiescent “+4” stem cells, although subsequent studies have shown that actively cycling crypt base stem cells also express these markers at equivalent levels^54^. Similarly, we did not detect enhanced expression of these markers in qCSCs. This is also in agreement with a recent single-cell RNA-sequencing analyses of the regenerating mouse intestine that identified a damage-induced quiescent cell type termed revival stem cells (revSCs)^15^. These cells, required for the regeneration of a functional intestine, are extremely rare during normal homeostasis and are characterised by enhanced expression of the pro-survival stress response gene Clu^55^. Interestingly, we find that many of the genes that make up the molecular signature of these quiescent revSCs are also enriched in qCSCs and have been found to regulate therapy resistance in various types of cancer. These common genes include *CLU*^56^, *CTSD*^57,58^, *CDKN1A*^59–63^, *EMP1*^64,65^, *MUC3*^66^, *LAMC2*^67^, *KRT19*^68^, *LGALS3*^69^, *F3*, *ITM2B*, *ITGB4*^70,71^, *CDH17*^72,73^ and *GSN*^74,75^ (Figure 3E – F, Figure S1 and Supplementary Data File 1). Considering that colon cancer is a heterogeneous tumor that recapitulates the cellular hierarchy of the intestine, these data suggest that the qCSCs identified here may be the tumor equivalent of revSCs. However, in contrast to revSCs and previous studies on qCSCs, our data demonstrate that qCSCs are enriched for p53 signaling (Figure 3A).

### qCSCs are dependent on p53 signaling

Loss of p53 in hematopoietic (HSCs) and neural stem cells (NSCs) causes these cells to exit quiescence and enter the cell cycle^76–78^. Targeting p53 may have the same effect in qCSCs but is complicated by the role of p53 as a tumor suppressor and guardian of homeostasis^79^. However, targeting negative cell cycle regulators downstream of p53 may provide novel strategies for qCSC elimination without affecting the role of p53 in healthy cells. Differential gene expression analysis, comparing PKH26^Negative^ and PKH26^Positive^ cells, identified the negative cell-cycle regulators *AKAP12*^80–83^*, CD82*^84^*, CDKN1A*^85–88^*, FHL2*^89–92^*, GPX3*^93–95^*, KIAA0247*^96,97^*, LCN2*^98–100^*, TFF2*^101–105^*, UNC5B*^106–108^ and *ZMAT3*^109,110^ to be enriched in qCSCs (Figure 3D). Significantly, each of these genes is a target of p53^79,80,82,92,96,109,111–115^, and with the exceptions of LCN2 and *ZMAT*3, associated with reduced survival in CRC (Figure 4A). Interestingly, CD82, KIAA0247 and UNC5B proteins localize to the cell surface and may therefore have potential as new markers for the prospective isolation of qCSCs in CRC. Indeed, CD82 has previously been identified as a marker for prospectively isolating stem cells from human fetal and adult skeletal muscle and is a functional surface marker of long-term HSCs^84,116^.

**Figure 4.**
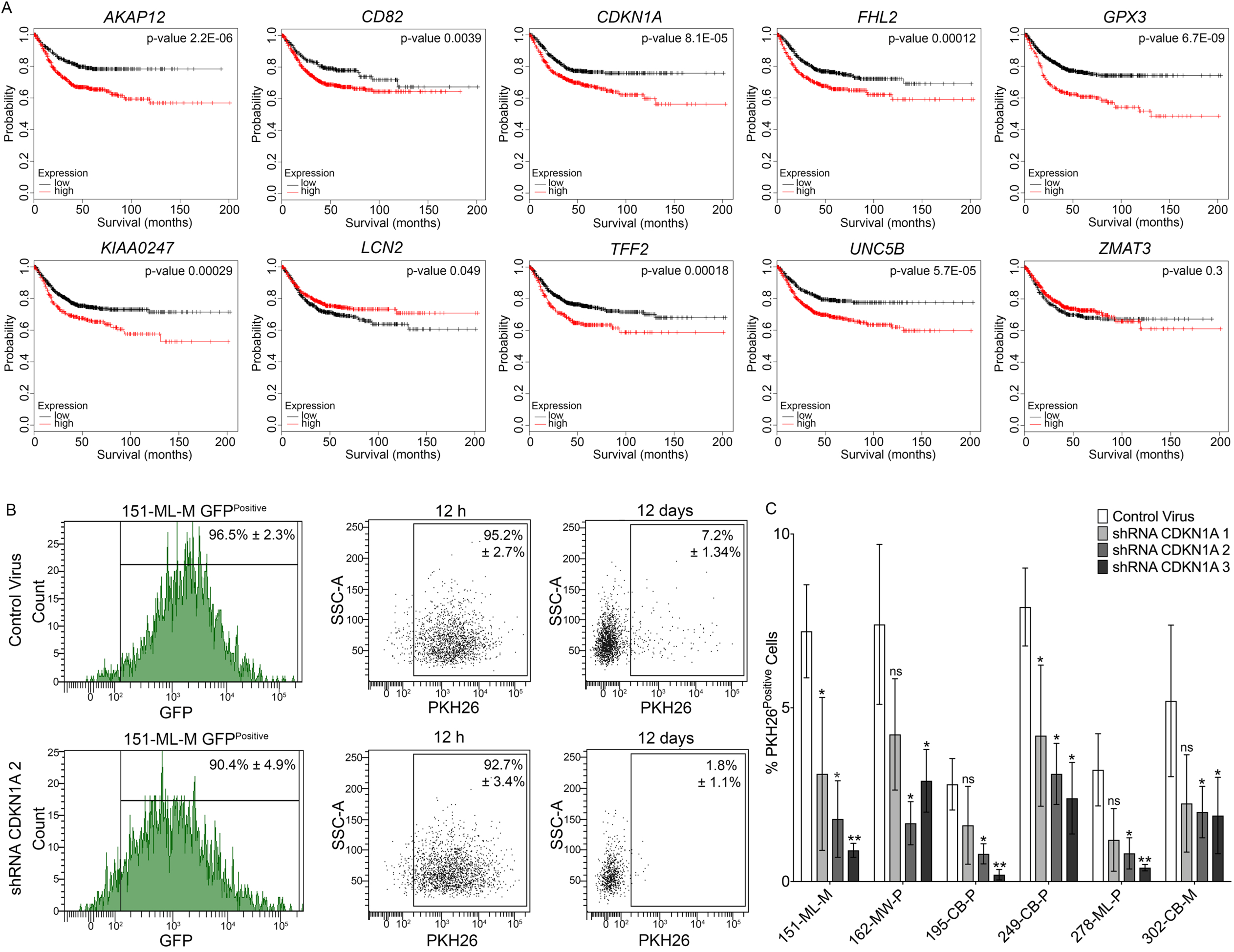
p53 target genes are indicators of poor prognosis and required for the maintenance of PKH26^Positive^ quiescent CSCs. (A) Kaplan-Meier survival curves for *AKAP12, CD82, CDKN1A, FHL2, GPX3, KIAA0247, LCN2, TFF2, UNC5B* and *ZMAT3* in colorectal cancer patients comparing lower quartile to upper quartile (logrank p-values = 2.2E-06, 0.004, 8.1E-05, 0.00012, 0.0003, 0.049, 0.00018, 5.7E-05 and 0.3, respectively). Of these, higher *AKAP12, CD82, CDKN1A, FHL2, GPX3, KIAA0247, LCN2, TFF2 and UNC5B* are significant at FDR < 10%. Results based upon data generated by the Kaplan-Meier Plotter (kmplot.com)^173^. (B) Representative FACS plot of PKH26 labelled 151-ML-M Control-GFP cells (top row) and shRNA CDKN1A-GFP cells (bottom row) after 12 h and 12 days. (C) Frequency (±SD) of PKH26^Positive^ LRCs in shRNA CDKN1A PDO models after 12 days compared to control virus transduced cells (data from 3 independent experiments). (See also Figure S2).

Deletion of *CDKN1A* (P21), which is the downstream mediator of p53 induced cell cycle arrest^86,117^, leads to cell cycle activation and exhaustion of quiescent HSCs and NSCs^118,119^. In addition, *CDKN1A* is highly expressed in noncycling intestinal crypt base stem cells^52^ and revSCs^15^. We therefore selected *CDKN1A* as a candidate gene to determine whether targeting the p53 pathway would eliminate qCSCs. Significantly, shRNA mediated knockdown of *CDKN1A* (Figure S3) in PKH26 labelled qCSCs resulted in the elimination of PKH26^Positive^ label-retaining qCSCs (Figure 4B and C).

## DISCUSSION

Colon cancer is a heterogeneous tumor entity containing a subpopulation of qCSCs that may promote tumor cell heterogeneity, plasticity, and resistance to various types of stress, including resistance to conventional treatments^29^. However, the rarity and plasticity of qCSCs has made them an elusive and challenging cell state to define and target. Here, we provide the first whole-transcriptome analyses of a population of colon cancer patient-derived long-term label-retaining qCSCs and identify genes that may provide novel targets for their elimination.

Label retention has previously been used as a strategy for the isolation of both healthy quiescent tissue stem cells and qCSCs from a variety of cancer types^11,17,25,26,29,120–122^. In agreement with these studies, we show that PKH26^Positive^ LRCs isolated from colon cancer PDOs are qCSCs capable of entering the cell cycle and self-renewing after replating in adherent and non-adherent cell culture conditions and maintain long-term quiescence in xenograft models. Interestingly, *in vivo* these cells were located at the tumor border, suggesting that quiescence may be induced at the invasive tumor front where such cells may be primed for metastatic dissemination. This is in agreement with previous studies showing cell cycle arrest / decreased proliferation and increased levels of Wnt signaling at the invasive front of colorectal tumors^123–127^.

RNA-sequencing of qCSCs demonstrated that they display the molecular hallmarks of quiescence^128^ while also being enriched for the same developmental and stem cell associated gene sets previously described for actively cycling ALDH^Positive^ CSCs^44^, which unlike PKH26^Positive^ LRCs are enriched in PDOs.

We previously reported that hedgehog signaling in active colon CSCs is non-canonical (SHH-dependent, PTCH-dependent, SMO-independent, GLI-independent) and acts as a positive regulator of Wnt signaling for CSC survival^44^. In agreement with our work, a subsequent study from Buczacki *et al*. (2018) demonstrated that qCSC survival in CRC is also dependent on non-canonical hedgehog signaling mediated regulation of Wnt signaling^37^. In addition, several of the genes common to both the revSCs reported by Ayyaz *et al* (2019)^15^ and qCSCs, namely CLU^129^, CTSD^130^, CDKN1A^131^, EMP1^132^, MUC3^133^, LAMC2^134^, KRT19^135^, LGALS3^136^, *F3*^137,138^, ITGB4^139^, CDH17^140^ and GSN^141^, are targets and/or regulators of Wnt signaling. Overall, these data demonstrate that both cycling and non-cycling CSCs share overlapping molecular profiles and further support the targeting of non-canonical hedgehog signaling to prevent disease relapse^37,44,142^.

However, the molecular mechanisms that distinguish non-cycling qCSCs from cycling CSCs required further elucidation. p53 plays a crucial role in regulating cellular stress responses such as DNA-damage repair, senescence, apoptosis and cell cycle arrest in virtually all cell types^87,143^. In addition, it is an important regulator of stem cell self-renewal and differentiation in embryonic and adult tissue stem cells^77,144,145^ and cancer stem cells^146–148^. Significantly, it has also been demonstrated to be essential for the maintenance of quiescence in HSCs, NSCs, muscle stem cells and lung progenitor cells^76,78,149–151^.

Here we show that qCSCs, in contrast to cycling ALDH^Positive^ CSCs^44^, are enriched for p53 signaling genes. p53 is mutated in 40 - 50% of CRCs. Reflecting this, half the tumors included in our study contain a p53 mutation (Table S1). However, regardless of mutation status, p53 appears to be functional in all the PDO models analyzed, as observed by p53-dependent expression of *CDKN1A* (Figure 3D)^152,153^.

Inhibiting the p53 pathway may therefore provide novel therapeutic “lock-out” strategies to induce the proliferation of qCSCs and thereby sensitize them to chemotherapeutics and prevent relapse^128,154,155^. Considering the role of p53 as a tumor suppressor and guardian of homeostasis in healthy tissues, as well as its inactivation in many cancers, most strategies to date have focused on the development of p53 activators^156^. However, our data, and others, suggest that strategies that activate p53 may lead to therapy resistance. For example, in breast cancer p53 induces senescence, drives resistance to therapy and is associated with poor therapeutic response and overall survival^157,158^.

Inhibiting p53 could interfere with its role in normal tissue homeostasis or lead to the activation of senescent cancer cells in other tissues. However, healthy cells have lower p53 expression levels than cancer cells^159^ and single dose treatments, that avoid the unwanted consequences of sustained p53 inhibition, may be sufficient to eliminate qCSCs. This was recently demonstrated by Webster *et al.* (2020) in melanoma, where a single dose of p53 inhibitor during the early stage of BRAF/MEK inhibitor treatment resulted in improved response to therapy^160^.

In addition, targeting negative cell cycle regulators downstream of p53, such as those identified here (*AKAP12*^80–83^*, CD82*^84,112^*, CDKN1A*^85–88^*, FHL2*^89–92^*, GPX3*^93–95^*, KIAA0247*^96,97^*, LCN2*^98–100,115^*, TFF2*^101–105^*, UNC5B*^106–108^ and *ZMAT3*^109,110^), may provide novel strategies for activating cell cycle in qCSCs without affecting the role of p53 in healthy cells. For example, p53-dependent activation of p21 (*CDKN1A*), which we show is required for the maintenance of qCSCs, is an important axis in senescence-dependent tumor suppression. However, despite p21 playing an important role in mediating the p53-dependent cellular response to stress, lack of p21 does not promote tumor development^161^. Furthermore, p21 maintains CSC self-renewal, limits proliferation and confers therapy resistance in numerous cancers types in which its temporal inhibition has been proposed as a strategy to overcome resistance to DNA-damaging chemotherapy and radiation^60–63,162–166^. Indeed, several small molecule inhibitors of p21 have been reported, including butyrolactone I^167^, LLW10^168^, sorafenib^169^ and UC2288^170^, that could serve as novel drugs for the elimination of therapy resistant qCSCs.

These data demonstrate the existence of long-term p53-dependent qCSCs in colon cancer and provide evidence supporting the temporal inhibition of p53 signaling, in combination with standard-of-care treatments, for the elimination of qCSCs and prevention of disease relapse. The p53 target genes identified here, along with the publication of our qCSC whole-transcriptome data, will provide a valuable resource for the development of such therapeutic strategies in the future.

## EXPERIMENTAL PROCEDURES

### Human tissue samples and establishment of patient-derived cancer organoid cell cultures

Tumor material was obtained with informed consent from CRC patients under approval from the local Institutional Review Board of Charité University Medicine (Charité Ethics Cie: Charitéplatz 1, 10117 Berlin, Germany) (EA 1/069/11) and the ethics committee of the Medical University of Graz and the ethics committee of the St John of God Hospital Graz (23-015 ex 10/11). Tumor staging was carried out by experienced and board-certified pathologists (Table S1). Cancer organoid cultures were established and propagated as described^45,171^.

### Cell cycle analysis and colony forming assays

Cell cycle analysis was carried using the Click-iT EdU assay (Invitrogen, #C10337) and assessed by FACS on a BD LSR II analyzer. For colony forming assays, PDOs were processed to single cells and labelled with CellTracker™ CM-DiI fluorescent dye (C7000, Thermo Fisher) or PKH26 (PKH26GL, Sigma-Aldrich) following manufacturer’s instructions and DAPI (to exclude dead cells). PKH26^Positive^ DAPI^Negative^ (live) cells were sorted by FACS (BD FACS Aria II) into adherent Matrigel culture. After 12 days, PDOs were once again processed to single cells and sorted by FACS, seeding PKH26^Positive^ DAPI^Negative^ cells and PKH26^Negative^ DAPI^Negative^ cells separately at limiting dilution into 96-well adherent Matrigel and 384-well non-adherent ultra-low attachment plates at a frequency of 100 and 1 cell per well, respectively. The purity of the sorted PKH26^Positive^ cell population was confirmed by post-sort FACS analysis. PDO sizes were determined by ImageJ software analysis. Ultra-low attachment wells containing spheroids were counted and used to calculate the CSC frequency using ELDA software (http://bioinf.wehi.edu.au/software/elda/index.html; Hu and Smyth, 2009).

### Xenotransplantation

Housing and handling of animals followed European and German Guidelines for Laboratory Animal Welfare. Animal experiments were conducted in accordance with animal welfare law, approved by local authorities, and in accordance with the ethical guidelines of Bayer AG. PDOs were processed to single cells and labelled with PKH26 (PKH26GL, Sigma-Aldrich) following manufacturer’s instructions and DAPI (to exclude dead cells). PKH26^Positive^ DAPI^Negative^ cells were collected by FACS and immediately transplanted by injected subcutaneously in PBS and Matrigel (1:1 ratio) into female 8 – 10-week-old nude^−/−^ mice at 1000 cells per animal. The purity of the sorted PKH26^Positive^ cell population was confirmed by post-sort FACS analysis.

### Immunofluorescence staining

Tumors were fixed in 4% paraformaldehyde overnight and cryopreserved in OCT compound. Immunohistochemistry of frozen sections was carried out via standard techniques with α-Tubulin (T5168, mouse monoclonal, Sigma; diluted 1:1000) and a secondary conjugated antibody at room temperature for 2 hours. For immunofluorescence imaging of PDOs, cultures were fixed in 4% paraformaldehyde for 30 min at room temperature and permeabilized with 0.1% Triton X-100 for 30 min and blocked in phosphate-buffered saline (PBS) with 10% bovine serum albumin (BSA). F-actin was stained with Alexa Fluor^®^ 647 Phalloidin (#A22287, Thermo Fisher; diluted 1:20) for 30 min at room temperature. Nuclei were counterstained with DAPI. Negative controls were performed using the same protocol with substitution of the primary antibody with IgG-matched controls. Cancer organoids were then transferred to microscope slides for examination using a Zeiss LSM 700 Laser Scanning Microscope.

### RNA Sequencing

Cells were lysed in RLT buffer and processed for RNA using the RNeasy Mini Plus RNA extraction kit (Qiagen). Samples were processed using NuGEN’s Ovation RNA-Seq System V2 and Ultralow V2 Library System and sequenced on an Illumina HiSeq 2500 machine as 2×125nt paired-end reads. The raw data in Fastq format were checked for sample quality using our internal NGS QC pipeline. Reads were mapped to the human reference genome (assembly hg19) using the STAR aligner (version 2.4.2a). Total read counts per gene were computed using the program “featureCounts” (version 1.4.6-p2) in the “subread” package, with the gene annotation taken from Gencode (version 19). Variance-stabilising transformation from the Bioconductor package DESeq2^172^ was used for normalisation and differential-expression analysis.

### Viral transduction

Cells were seeded in 100 μl volumes of antibiotic free culture media at 1.0 x10^5^ cells per well in ultra-low attachment 96-well plates. Control and shRNA lentiviruses were purchased from Sigma-Aldrich (Table S2). Viral particles were added at a multiplicity of infection of 1. Cells were transduced for up to 96 h or until GFP positive cells were observed before being embedded in Matrigel for the establishment of lentiviral transduced cancer organoid cultures. Puromycin (2 μg/ml) was used to keep the cells under selection.

### Gene expression analysis

For quantitative real-time RT-PCR analysis RNA was isolated using the RNeasy Mini Plus RNA extraction kit (Qiagen). cDNA synthesis was carried out using a Sensiscript RT kit (Qiagen). RNA was transcribed into cDNA using an oligo dTn primer (Promega) per reaction. Gene expression analysis was performed using TaqMan^®^ Gene Expression Assays (Applied Biosystems) (Table S3) on an ABI Prism 7900HT sequence detection system (Applied Biosystems). GAPDH was used as an endogenous control and results were calculated using the Δ-ΔCt method. Data were expressed as the mean fold gene expression difference in three independently isolated cell preparations over a comparator sample with 95% confidence intervals. Survival curves were generated using the Kaplan-Meier Plotter (www.kmplot.com/analysis)^173^. Gene ontology enrichment analysis was carried out using the Gene Ontology Resource (www.geneontology.org)^174,175^.

### Statistical analysis

GraphPad Prism 8.0 was used for data analysis and imaging. All data are presented as the means ± SD, followed by determining significant differences using the two-tailed t test. Significance of RT-PCR data was determined by inspection of error bars as described by Cumming *et al*. (2007)^176^. Gene set enrichment analysis was carried out using pre-ranked feature of the Broad Institute GSEA software version 2 using msigdb v5.1 gene sets^177,178^. The ranking list was derived from the fold changes calculated from the differential gene expression calculation and nominal p-values. P-values <0.05 were considered as statistically significant. The representation factor and the associated probability of finding an overlap were calculated using http://nemates.org/MA/progs/representation.stats.html. Survival curves were generated using the Kaplan-Meier Plotter (www.kmplot.com/analysis)^173^. For the final list of significant genes, False Discovery Rate (FDR) was computed using the Benjamini-Hochberg method^179^.

## Acknowledgements

We thank Dorothea Przybilla, and Cathrin Davies (Laboratory of Molecular Tumor Pathology, Charité Universitätsmedizin Berlin, Germany) for technical and cell culture assistance. The research leading to these results has received support from the Innovative Medicines Initiative Joint Undertaking under Grant Agreement 115234 (OncoTrack), the resources of which are composed of financial contribution from the European Union Seventh Framework Programme (FP7/2007-2013) and EFPIA companies in kind contribution. A.S., T.J., D.M. and. D.H. are employees of Bayer AG. R.L., J.T. and M.L. are employees of Nuvisan ICB GmbH. C.R.A.R.

## Authors Contribution

Conceptualization, J.L.R.; Methodology, J.L.R.; Investigation, J.L.R., D.S., S.S., A.S., R.L., J.T., T.J., J.H., and M.L.; Writing, J.L.R.; Visualization, J.L.R; Data Curation, A.S., J.T.; Resources, J.H., U.K., C.R.A.R. and B.G.; Supervision, J.L.R., D.M., D.H., R.S., and M.L.

## Accession Numbers

Array data are available in the ArrayExpress database (www.ebi.ac.uk/arrayexpress) under accession number E-MTAB-8924.

## Supplemental Information

**Figure S1:**
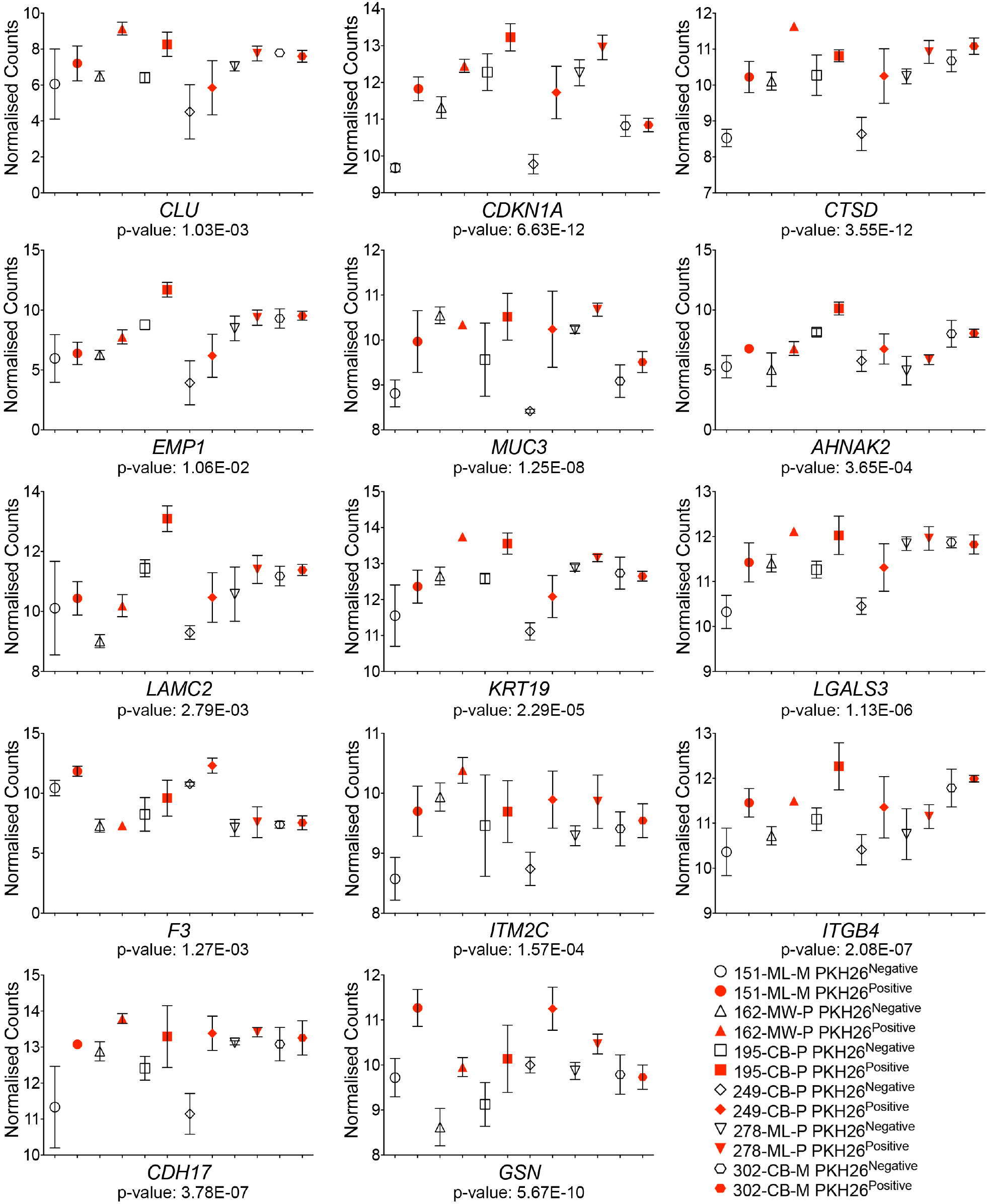
RNA-sequencing generated normalized counts for differentially expressed and common revSC^1^ molecular signature genes in PKH26^Positive^ qCSCs compared to PKH26^Negative^ cells. Related to Figure 3.

**Figure S2:**
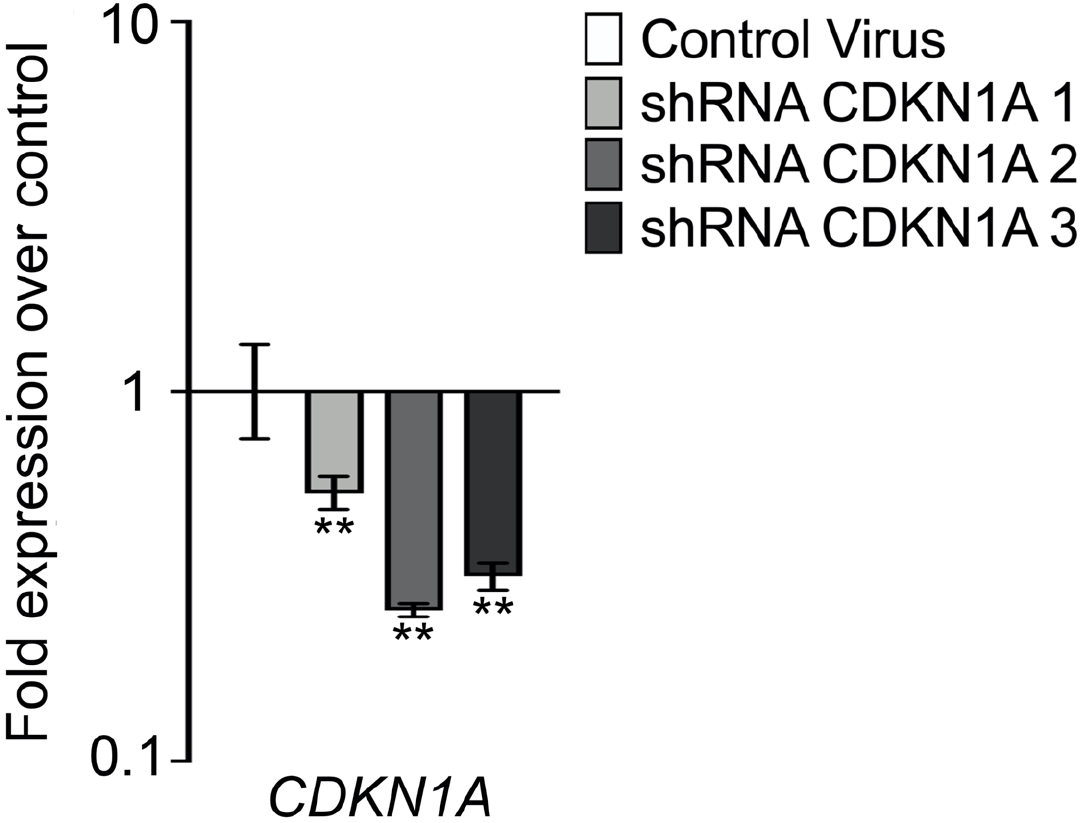
Fold expression of CDKN1A (±95% confidence intervals) in shRNA CDKN1A transduced PDOs from three independent experiments. Related to Figure 4. Significant differences are *p-value < 0.05; **p-value < 0.01 and were determined by inspection of error bars as described by Cumming et al. (2007)^2^.

**Table S1.** Tissue Origin, TNM Classification and P53 status of tumors. Related to Figure 1. T: primary tumor size, N: regional lymph nodes involved, M: distant metastasis, *SNV: non-synonymous single nucleotide variant

| Patient Model | Origin | TNM stage | Stage | P53 Status |
| --- | --- | --- | --- | --- |
| 151-ML-M | Liver | T2 N0 M0 , M1a | IVA | *SNV (G266E) |
| 162-MW-P | Sigmoid colon & descending colon | T3 N0 M0 | IIA | Wild type |
| 195-CB-P | Sigmoid colon | T4a N2b M1a | IVA | *SNV (C135F) |
| 249-CB-P | Ascending colon | T3 N0 M0 | III | Wild type |
| 278-ML-P | Sigmoid colon & descending colon | T4a N0 M0 | IIB | *SNV (R273C) |
| 302-CB-M | Liver | T3 N1a M1a | IVA | Wild type |

**Table S2.**
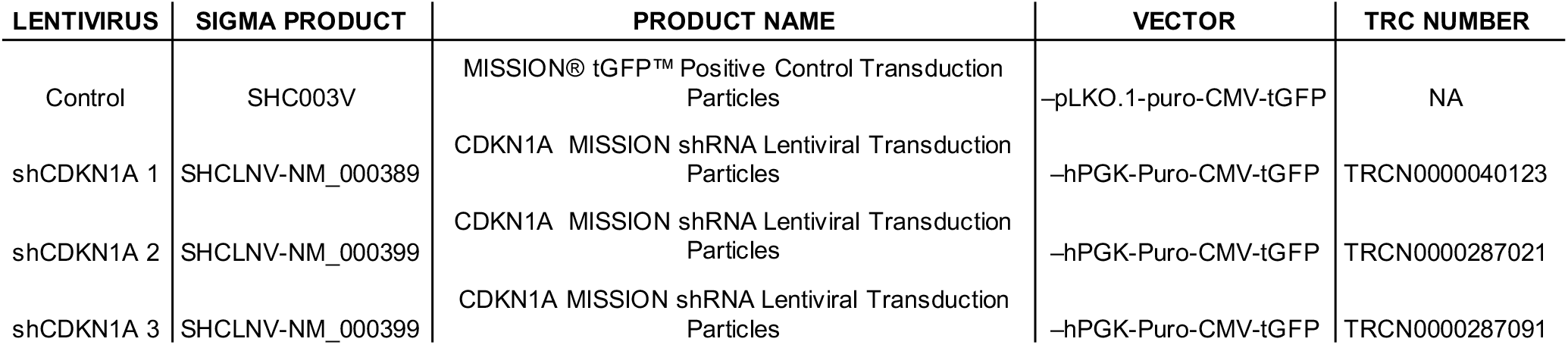
Lentiviral Transduction Particles. Related to Figure 4.

| LENTIVIRUS | SIGMA PRODUCT | PRODUCT NAME | VECTOR | TRC NUMBER |
| --- | --- | --- | --- | --- |
| Control | SHC003V | MISSION® tGFP™ Positive Control Transduction Particles | -pLKO.1-puro-CMV-tGFP | NA |
| shCDKN1A 1 | SHCLNV-NM_000389 | CDKN1A MISSION shRNA Lentiviral Transduction Particles | -hPGK-Puro-CMV-tGFP | TRCN0000040123 |
| shCDKN1A 2 | SHCLNV-NM_000399 | CDKN1A MISSION shRNA Lentiviral Transduction Particles | -hPGK-Puro-CMV-tGFP | TRCN0000287021 |
| shCDKN1A 3 | SHCLNV-NM_000399 | CDKN1A MISSION shRNA Lentiviral Transduction Particles | -hPGK-Puro-CMV-tGFP | TRCN0000287091 |

**Table S3.** Taqman^®^ Gene Expression Assays. Related to Figure 4.

| Symbol | Gene Name | UniGene ID | TaqMan® Gene Expression Assay |
| --- | --- | --- | --- |
| CDKN1A | cyclin dependent kinase inhibitor 1A | Hs.370771 | Hs00355782_m1 |
| GAPDH | glyceraldehyde-3-phosphate dehydrogenase | Hs.544577 | Hs02758991_g1 |

